# Hyperglycemia worsens smooth muscle foam cell formation through PI3Kγ-dependent defective autophagy

**DOI:** 10.1101/2025.01.21.634036

**Authors:** Labrana Honora, Wahart Amandine, Cormier Kévin, Solinhac Romain, Swiader Audrey, Mentouri Isra, Smirnova Natalia, Malet Nicole, Gayral Stéphanie, Ramel Damien, Auge Nathalie, Laffargue Muriel

**Author notes:** Authors for correspondence: Muriel Laffargue, PhD or Nathalie Augé, PhD., or. equally contributed (Labrana Honora, Wahart Amandine, Auge Nathalie, Laffargue Muriel).

## Abstract

**Background:** Diabetes significantly increases the risk of cardiovascular complications, particularly through the mechanisms of atherosclerosis. In this context, uncontrolled hyperglycaemia is a key contributor to arterial dysfunctions. However, the specific effects of elevated glucose levels on smooth muscle cells (SMC)-derived foam cells formation remain poorly defined.

**Methods:** Using a wire injury-based accelerated atherosclerosis mouse model combined to lipid biochemistry and cellular lipid imaging approaches, we analysed molecular mechanisms involved in SMC derived foam cell formation.

**Results:** Our findings demonstrated that hyperglycaemia induced a specific lipid loading in SMC in an accelerated atherosclerosis mice model. *In vitro*, high glucose concentration negatively affected autophagic process independently of lipid uptake and efflux in human and mouse primary SMC. Consistently, treatment with autophagy activators successfully reduced lipid droplet formation. Mechanistically, we identified that PI3Kγ activity impaired autophagy through TFEB phosphorylation promoting SMC foam cells formation in response to high glucose.

**Conclusion:** These findings shed light on the mechanisms underlying SMC homeostasis disruption under hyperglycemic conditions and underscore the potential therapeutic window for PI3Kγ inhibitor in managing cardiovascular diseases in diabetic patients.

## INTRODUCTION

Atherosclerosis is a chronic inflammatory disease that begins with the infiltration of lipids into the arterial intima and culminates with the formation of thrombi, leading to stroke, infarction or ischemia of the lower limbs, depending on the location. The standard treatment for atherosclerosis involves performing angioplasty with stenting to restore arterial blood flow. Despite advancements in patient management, approximately 17.6 % of patients require revascularization within five years of their initial cardiovascular event ^1^. Diabetic patients represent a population at particularly high risk for recurrence ^2^. Atherosclerosis is notably accelerated in these individuals, leading to the rapid development of complicated plaques. The factors incriminated in this acceleration are dyslipidemia, characterized by elevated levels of atherogenic low-density lipoproteins (LDL), hyperglycemia, oxidative stress and increased inflammation. In particular, hyperglycemia has been proven to exert deleterious effects on endothelial cells, immune cells and smooth muscle cells (SMCs), which are major contributors to the development of atherosclerosis ^3^. It has been put forth that hyperglycemia precipitates the dedifferentiation of SMCs ^4^. In their dedifferentiated state, SMCs constitute the predominant cell type in early arterial intimal thickenings and represent an important component of most stages of atherosclerosis ^5^. Their remarkable plasticity gives them the ability to take on different phenotypes. Recent single-cell studies have revealed the presence of several subpopulations of synthetic SMCs within human and mouse atherosclerotic plaques, including fibroblast-like cells, chondrocyte-like cells, and cells capable of phagocytosing lipids. These SMC-derived cell phenotypes participate together with macrophages to the pool of intra- plaque foam cells ^6–8^. Although the proportion of foam cells originating from SMCs *versus* macrophages remains controversial, the existence of these smooth muscle foam cells and their role in atherosclerosis are currently well accepted. Whereas the role of hyperglycemia has been largely studied in macrophages (Qi et al., 2023), whether or not this hyperglycemia could impact foam SMCs has not been clearly investigated.

In order to respond to this significant enquiry, we are utilizing an *in vivo* murine model of hyperglycemia and an *in vitro* model of SMCs exposed to low or high concentrations of glucose. *In vivo*, we demonstrated that hyperglycemia has a specific impact on the SMC lipid loading in intimal hyperplasia induced by an endovascular injury in Western diet-fed LDLR-/- mice. This finding was validated *in vitro* in primary SMCs incubated in high glucose conditions. Furthermore, our findings revealed that this effect was attributable to a reduction in autophagy, induced by high concentrations of glucose and could be reversed by autophagy activators. We then demonstrated an unanticipated role of the specific PI3Kγ isoform in these processes through the regulation of TFEB, a transcription factor governing autophagy. Finally, inhibiting PI3Kγ proved effective in restoring a normal autophagic flux and blocking the development of foam SMCs.

## MATERIALS AND METHODS

### Availability of Data and Materials

Upon reasonable request to the corresponding authors, the data and materials supporting the findings of this study are available.

### Antibodies

The rabbit anti-α-SMA polyclonal antibody (#RB9010P1) was purchased from Epredia. The rabbit anti-LC3 polyclonal antibody (#2775), the rabbit phospho-TFEB (S211) monoclonal antibody (#37681), the rabbit anti-GAPDH monoclonal antibody (#2118), the anti-mouse IgG HRP-linked (#7076), and the anti-rabbit HRP-linked antibodies (#7074) were purchased at Cell Signaling Technology. Rabbit polyclonal TFEB antibody used for Western Blot analysis (#13372-1-AP) was bought at Bethyl Laboratories and Proteintech respectively. The mouse anti-histone deacetylase 1 (HDAC1) monoclonal antibody (sc-8410) was purchased at Santa Cruz Biotechnology, Inc. The immunofluorescence analysis secondary antibodies AlexaFluor488 goat anti-rabbit (#A11008), and AlexaFluor555 goat anti-rabbit (#A32732) were purchased at Invitrogen.

### Other materials and Reagents

Dulbecco’s Modified Eagle Medium, no glucose (DMEM, 11966-025), Fœtal Bovine Serum (FBS, A31608-01) and Opti-MEM (31985-047) were purchased from Gibco. D-Glucose (G7021), Streptozotocin (S0130), Rapamycin (R0395), LY294002 (L9908), XtremeGENE 9 DNA (6365787001), Nile Red (N-3013) DAPI (D9542) and Bafilomycin (B1793) were purchased from Sigma-Aldrich and NBD-cholesterol (N1148) from Life Technologies. BSA (04- 100-812C) was from Euromedex and 3-MethylAdenine (HY-19312) from Clinisciences. A66 (S2636), AZD 6482 (S1462) and IPI-549 (S8330) were obtained from SelleckChem. DilC18 1,1’dioctadecyl-3,3,3’,3’tetrametyl-indocarbocyanine perchlorate (D3911), Lipofectamine 2000 (H668-019) and Bodipy 493/503 (D3922) were purchased from Invitrogen.

### Animals studies

Male C57Bl/6J LDL receptor-deficient (LDLR^−/−^) mice and male Wild-Type C57BL/6J (WT) mice were obtained from Charles River Laboratories (Wilmington, MA, USA) and ENVIGO (Gannat) respectively. Male PI3Kγ-deficient (PI3Kγ KO) mice were from a C57BL/6J background and have been described earlier ^9^. All mice were maintained in specific opportunist and pathogen-free conditions at the CREFRE breeding facility (US006, Toulouse, France) and used in accordance with institutional guidelines from Directive 2010/63/EU of the European Parliament or the NIH guidelines for the protection of animals used for scientific purposes.

For the injury-induced atheromatous protocol, eight-weeks-old LDLR^−/−^ male mice were fed with Western diet (42% from fat, 0,2% cholesterol, MD.88137, ENVIGO) for 9 weeks. After 2 weeks of dietary adaptation, mice were randomly assigned to receive low doses of streptozotocin (50 mg/kg, STZ group) or vehicle (Control group) for 5 days. Then, mice underwent carotid artery surgery and were euthanized 4 weeks later for blood and tissue collection.

### Carotid Artery Wire Injury

Endovascular injury of the carotid artery was performed as previously described ^10^. Briefly, mice were anaesthetized with an intraperitoneal injection of a ketamine (100 mg/kg)/xylazine (20 mg/kg) mix followed by a buprenorphine injection (100 µg/kg). The left common carotid artery was isolated and the external carotid incised under a surgical stereomicroscope (Carl Zeiss). A 0.35-mm diameter angioplasty guidewire (Abbott Vascular) was introduced into the artery and withdrawn three times (5 mm length in total) to induce de-endothelialization. The external carotid artery was ligated, post-wire removal, restoring blood flow to the common and left carotid artery. After 4 weeks, the injured carotid arteries were collected for subsequent immunofluorescence analysis.

### Tissue Processing and Lesion Analysis

Injured carotids were frozen on a cryostat mount with optimum cutting temperature (n°4583; Tissue-Teck) and sectioned into 10 µm segments from the common carotid over the length of 250 µm using a cryostat microtome (Cryostar NX50, Microm Microtech). Sections were washed with PBS (Phosphate-Buffered Saline), fixed with 4% paraformaldehyde in PBS (PFA) and permeabilized with 0.1% of Triton X-100 in PBS and blocked with 3% bovine serum albumin (BSA) in 0.1% Triton in PBS for 1h. Subsequently, primary antibody was applied and incubated overnight at 4°C. Following secondary antibody incubation, neutral lipids were stained with Bodipy 493/503 (1:1000) for 30 min and nuclei were counterstained with DAPI (100µg/mL) for 5 min at room temperature (RT).

### Cell Culture

Human SMCs were isolated from mesenteric arteries of deceased anonymous donors ^11^. SMCs were isolated from three different mesenteries (anonymous N° 75224, 89049 and 92224) and were immortalized (hSMC) by transfection of SV40T antigen, using Lipofectamine LTX Reagent and Plus Reagent (Invitrogen/Life Technologies). Cells were routinely maintained in DMEM, 10% foetal bovine serum (FBS) supplemented with 1 g/L glucose, at 37°C in a humidified 5% CO2 air atmosphere. Experiments were realised with cells from passages 4 to 20. Mouse SMCs were isolated from WT and PI3Kγ KO mice according to a modified protocol described by Ray et al. ^12^. In brief, aortae from 10 WT and 10 PI3Kγ KO mice were dissected from their origin to the iliac bifurcation, flushed with PBS, and removed. The adventitia was removed from the aortae and the smooth tubes were cut into pieces of 1–2 mm, and then digested in 0.3% collagenase solution. Collagenase digestion was stopped by adding DMEM/10% FBS. Then, the pieces of aortae were washed twice and seeded in culture dishes coated with 10 µg/ml human fibronectin (BD Biosciences). Primary confluent cultures were trypsinized at 37°C, and then cells were incubated at 37°C in 5% CO2 in DMEM/10% FBS. Cells were cultured for 7 days in either low (1 g/L) or high (5 g/L) glucose medium to mimic normal and hyperglycemic conditions, respectively.

### si-RNA Transfection

SMCs were transfected using Lipofectamine 2000, according to the manufacturer’s protocol. si-RNA sequences were purchased at Qiagen. The two target sequences of si-RNA against Atg5 were as follows: TCCAACTTGTTTCACGCTATA (SI00069251) and AACCTTTGGCCTAAGAAGAAA (SI022655310). The two target sequences against Atg7 were: ATCAGTGGATCTAAATCTCAA (SI02655373) and ATCGGATGAATGAGCCTCCAA (SI04231360). Target sequences against TFEB were: GAGACGAAGGTTCAACATCAA (SI00094976) and CACAGGCTGTTGTGAGGACTA (SI04384240). For each experiment a comparison with a negative siRNA was performed (Cat. No.:1027281). The silencing efficiency was measured by Real-time PCR analysis.

### LDL Isolation and Oxidation

Human LDL and HDL (High-density lipoprotein) were isolated from pooled fresh sera by sequential ultracentrifugations, dialyzed and sterilized by filtration. LDL oxidation by UV-C irradiation was obtained as previously described ^13^. ApoA-1 was prepared from HDL as described before ^14^.

### Cellular Immunofluorescence

After stimulation, the cells plated onto coverslips were rinsed with PBS, fixed with 4% PFA, permeabilized for 10 min with PBS containing 0,1% Triton X-100 and blocked for 30 min in PBS 3% BSA. Cells were incubated, 1h with anti-LC3 primary antibody (1:100) and after rinsing, 1h with AlexaFluor488 goat anti-rabbit secondary antibody (1:500). Microscopy imaging was performed using a Zeiss LSM780 confocal microscope equipped with a Plan Apochromat 63× 1.4 NA oil-immersion objective lens. Images were analysed using ImageJ software.

### Autophagic Flux

Autophagic flux was measured using two techniques. SMCs were either treated with Bafilomycin (100 nM, Baf) 6 hours before the end of the experiment or were transfected with pcDNA3-GFP-LC3-RFP-LC3ΔG construct (Addgene, 168997 deposited by Noboru Mizushima) as described previously ^15^. Following the bafilomycin treatment LC3 expression was assessed using Western Blot analysis. LC3B-II flux was determined by subtracting the LC3B-II densitometric value from bafilomyin-untreated cells compared to bafilomycin-treated cells. Transfection was performed using XtremGENE 9 DNA transfection agent. 24 hours after transfection cells were washed and then fixed with 4% PFA in PBS for 15 min at RT. Nuclei were labelled with DAPI, 5 min at RT. Images were captured by LSM780 confocal microscope equipped with a Plan Apochromat 63× 1.4 NA oil-immersion objective lens. Fluorescence analysis was performed using ImageJ software and data from each condition were represented as the ratio of median GFP fluorescence to RFP fluorescence.

### Foam cell formation assay

SMCs lipid loading was quantified after cell fixation and permeabilization (as described for cellular immunofluorescence) by Nile Red (3 min) or Bodipy 493/503 (30 min) staining and DAPI counterstaining (1 µg/mL in PBS). Staining was assessed by fluorescence microscopy (ZOE™, Biorad) and quantified after fluorescence eluting with NaOH solution (0,3 M) at 37°C (for Bodipy λex: 485 nm-λem:535nm/ for Nile red λex: 591nm-λem: 657nm, normalized to nucleus labelling). Alternatively, intracellular total and esterified cholesterol levels were quantified using a cholesterol fluorometric assay kit (Abcam, ab65359). The level of intracellular cholesterol was normalized to the protein concentration of the sample.

### Di-oxidized LDL uptake assay

SMCs cultured in 1 g/L and 5 g/L 10%FBS/DMEM medium for 7 days were passaged and incubated in 1 g/L and 5 g/L 1%FBS/DMEM medium, supplemented with carbocyanine-oxLDL (60 µg/mL) and oxLDL (100 µg/mL) for 24 hours (Di-oxL). After treatment, cells were fixed using 4% PFA in PBS, permeabilized with Triton X-100 0.1% in PBS and nuclei were counterstained with DAPI 5 min at RT. Images were captured using fluorescent microscopy (ZOE, Biorad) and then quantified after fluorescence eluting with NaOH solution (0,3 M) at 37°C (λex: 530nm-λem: 580nm normalized to nucleus labelling).

### Cholesterol efflux assay

After 7 days in 10%FBS/DMEM at 1 g/L or 5 g/L glucose, cells were passaged in 1%FBS/DMEM medium and loaded with 1 µg/mL NBD-cholesterol for 48 hours. Then, SMCs were rinsed with PBS and equilibrated in DMEM without phenol red/0,5% BSA medium for 2h, and incubated in 1 g/L and 5 g/L glucose 1% FBS/DMEM medium without phenol red containing ApoA-1 (100 µg/mL) and HDL (100 µg/mL) and 1 or 5 g/L glucose for 24 hours. Next, the medium was collected, and the cells were lysed at 37°C with 0.3 M NaOH solution. The fluorescence intensity was measured in both fraction with a TECAN microplate spectrophotometer (λex: 485 nm-λem:535nm). Efflux was calculated as follows: Efflux = NBDfluorescence in medium / (NBDfluorescence in medium + NBDfluorescence in cell lysate). Displayed values represent subtracting effluxes of the wells without apoA-I or HDL from those containing ApoA-I or HDL.

### Western Blot analysis

Cell extracts were prepared using RIPA lysis buffer supplemented with protease/phosphatase inhibitors. Nuclear/cytosolic extracts were obtained using the NE-PER Nuclear and cytoplasmic extraction reagents (#78835, Thermo Fisher Scientific) following manufacturer protocol. Protein levels were assessed by Bradford assay. For regular western, equal amount of protein samples was separated by SDS-PAGE. Nonspecific binding was blocked with 5% BSA in Tris buffer saline (TBS) for 2h at RT. Blots were incubated overnight at 4°C with following primary antibodies: anti-LC3 (1:1000), anti-TFEB (1 :1000), anti P-TFEB (S211) (1:1000). β-actin, GAPDH and HDAC1 were used as loading control. After 3 washes with Tris Buffered Saline (TBS) 0,1%Tween20, membranes were incubated with secondary antibodies for 1h at RT. Membranes were scanned using Chemidoc Touch imaging system (Biorad) and proteins were detected by ECL reagents. All the western blots were presented with protein ladder at the left of the membrane (see uncrop supplementary data). The bands intensities were quantified using ImageJ software.

### Real-time quantitative PCR

Total RNA was extracted from VSMCs using Trizol reagent and cDNA was amplified using a High-Capacity cDNA Reverse Transcription Kit (Thermo Fisher Scientific, 4368813) on a thermocycler (T100 Biorad #1861096). Real time quantitative PCR was performed using StepOnePlus Real Time PCR System and the results were analyzed using StepOne Software v2.3. The primers for the HPRT were 5ʹ- TTGCTCGAGATGTGATGAAGGA−3ʹ (forward) and 5ʹ- CCAGCAGGTCAGCAAAGAATT −3ʹ (reverse). The primers for human ABCA1 were 5’- CACCCCGTATGAACAGGATT-3’ (forward) and 5’- GCCAAGGACCAAAGTGATG-3’ (reverse). The primers for human ABCG1 were 5’-ATTCAGGGACCTTTCCTATTCGG-3’ (forward) and 5’- CTCACCACTATTGAACTTCCCG-3’ (reverse). The primers for the human LC3 receptor were 5ʹ- AGCAGCTTCCTGTTCTGGAT−3ʹ (forward) and 5ʹ- TGAGCTGTAAGCGCCTTCTAA −3ʹ (reverse). The primers for human LAMP-1 were 5ʹ- TCCAGGCGTACCTTTCCAACAG −3ʹ (forward) and 5ʹ-TGTCTTGTTCACAGCGTGTCTCTC −3ʹ (reverse). The primers for human Atg5 were 5ʹ- TGGATTTCGTTATATCCCCTTTAG−3ʹ (forward) and 5ʹ- CCTAGTGTGTGCAACTGTCCA −3ʹ (reverse). The primers for Atg7 were 5ʹ- CCGTGGAATTGATGGTATCTGT −3ʹ (forward) and 5ʹ-TCATCCGATCGTCACTGCT −3ʹ (reverse). The primers for human TFEB were 5ʹ- CCATCACCTGGACTTCAGCC−3ʹ (forward) and 5ʹ- CTTGGACAGGCTGGGGAATG −3ʹ (reverse). The primers for murine LC3 were 5ʹ- ATTGCTGTCCCGAATGTCTCʹ (forward) and 5ʹ- CGTCCTGGACAAGACCAAGT −3ʹ (reverse). The primers for murine LAMP-1 were 5ʹ- GCAACTTCAGCAAGGAAGAGAC−3ʹ (forward) and 5ʹ- AACATTGTACTTGGATACAGTGGG −3ʹ (reverse) and for murine HPRT CCTCCTCAGACCGCTTTTT (forward) and AACCTGGTTCATCATCGCTAA (reverse). The 2ΔΔCt method was used to calculate the fold changes in gene expression. HPRT served as the control for comparison. qRT-PCR was performed in triplicate and repeated in at least three separate experiments.

### Statistical Analysis

Data are presented as the mean ± standard error of measurement (SEM) of 3 to 8 independent experiments. The statistical analysis procedure was defined after performing a normality test (Shapiro-Wilk) to ensure equality of variances. To compare two experimental conditions, the results were analysed using Student’s parametric test or the non-parametric Wilcoxon or Mann Whitney tests, when the variances were equal or unequal respectively. Beyond 3 experiments, the data were compared using parametric two-factor ANOVA or non-parametric Friedman’s test, for equal or unequal variances respectively. Statistical analyses were performed using Prism 10 software (GraphPad software). Results were considered statistically significant when p<0.05.

## RESULTS

### High glucose concentration enhances lipid accumulation in smooth muscle cells *in vitro* and *in vivo*

To investigate the impact of hyperglycemia on atherosclerosic lesions, we developed a wire injury-based model of accelerated atherosclerosis in hyperglycemic mice. Briefly, an endovascular injury of the carotid artery was performed ^10^ in western diet-fed LDLR^-/-^ mice rendered hyperglycemic or normoglycemic respectively by repeated injections of low doses of streptozotocin (STZ) or vehicle (Ctrl) (Figure 1A). As expected, STZ-treated mice showed a significant increase in blood glucose levels after 7 days (325+/- 40.9 mg/dL *vs* 173.6+/- 9.7 mg/dL in Ctrl mice, ***p<0.001) and reached 514.5+/- 57.2 mg/dL at the end of the protocol (Table 1). One month after the carotid artery injury, both groups of mice (Ctrl and STZ) developed a lipid-rich neointima at the lesion site, mainly composed of SMCs as revealed by α-SMA labelling (Figure 1B-D). Despite a non-significant increase in the total lipidic area, lesions from STZ-treated mice exhibited a significant increase in colocalization of α-SMA with bodipy labelling (Pearson correlation coefficient) of 0.443+/-0.0375 *vs* 0.251+/-0.0.22 in ctrl mice, ** p< 0.01, Figure1D). These data suggest that STZ treatment led to increased foam SMC content in the neointima of atherosclerosis-prone injured arteries. Analysis of the blood metabolic profile of these mice revealed that despite a significant increase in plasmatic LDL, STZ-treated mice had an unchanged LDL/HDL ratio (Table 1). To further investigate the possible direct impact of hyperglycemia on foam SMC formation, we cultured human and mouse SMCs with low (L, 1 g/L) or high (H, 5 g/L) glucose concentration. After incubation of SMCs with oxidized low-density lipoprotein (OxL), we observed a dramatic increase in intracellular lipid accumulation in high glucose compared to low glucose condition in both human and mouse cells (Figure 1E, F respectively). Consistent with these changes, the total intracellular cholesterol, especially cholesterol ester rates, were also increased under elevated glucose concentration in human SMCs (Suppl. Fig. 1A, B). Altogether, our results demonstrate that high glucose concentration favours SMC-derived foam cell accumulation *in vivo* and *in vitro*.

**Figure 1:**
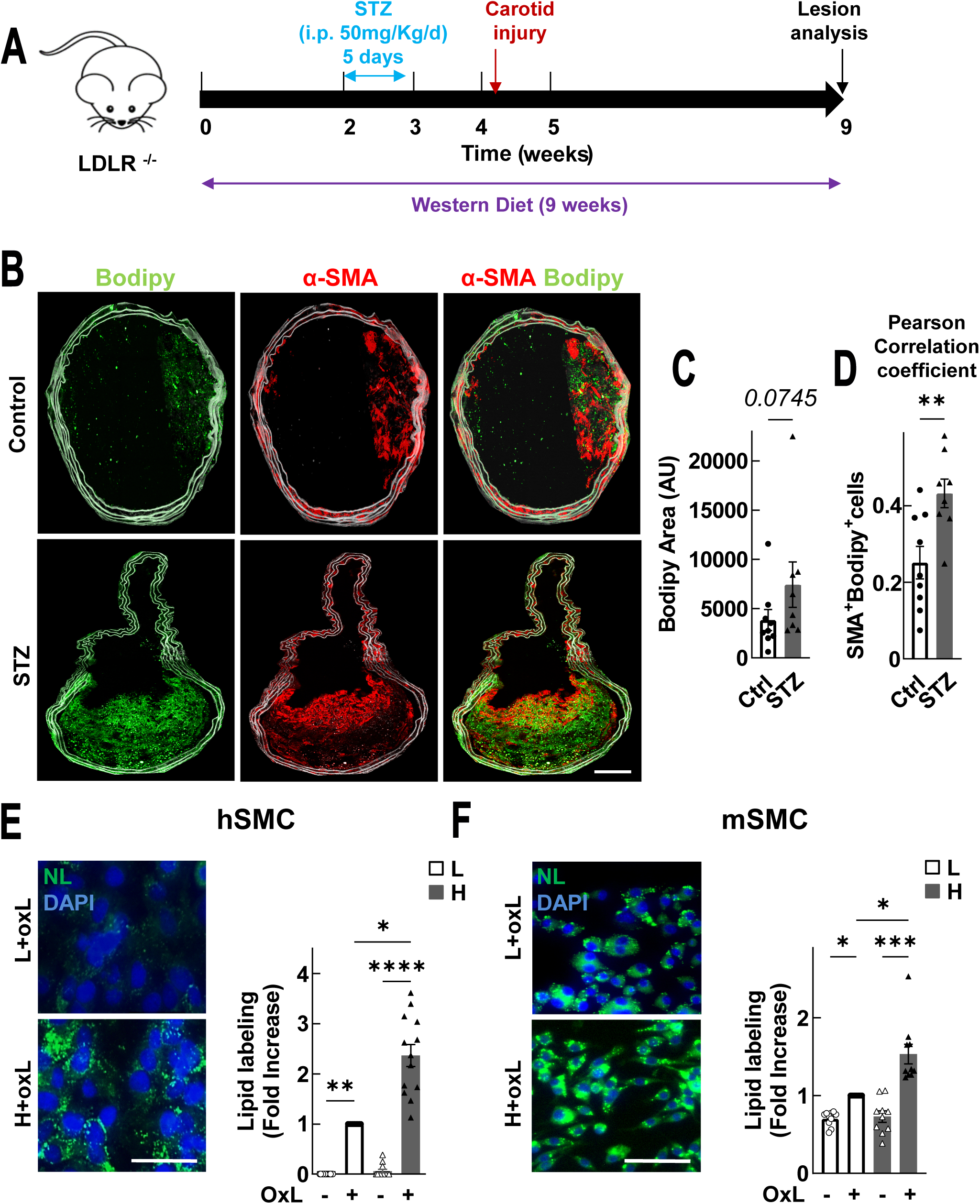
**High glucose concentration leads to increased lipid accumulation in smooth muscle cells *in vitro* and *in viv*o**: A: Summarized procedure to induce a hyperglycemic wire- injured atheromatous mouse model: LDLR^-/-^ mice fed with Western diet during 9 weeks, received 5 injections of either streptozotocin (STZ, 50 mg/kg, n=8) or citrate buffer vehicle (0.1 N, Ctrl, n=9). 10 days after the last STZ injection, mice underwent a carotid artery injury and were euthanized 4 weeks after. Blood and injured carotid arteries were harvested for analysis. B: Representative images of carotid artery sections stained with Bodipy 493/503 (green, neutral lipids) and α-SMA (red, smooth muscle cells, SMCs). Internal elastic lamina autofluorescence appears in grey. Scale bar: 100 µm. C: Neutral lipids inside the lesion were quantified by measuring the Bodipy labelling inside the neointima. D: Lipid accumulation in SMCs was quantified based on the merged red/green labelling. E, F: Representative images of neutral lipid accumulation (left) in human SMCs (hSMC, E) and murine SMCs (mSMC, F) incubated either in low (L, 1 g/L) or high (H, 5 g/L) glucose medium for 7 days, followed by oxLDL (oxL, 200 µg/mL) exposure for 3 additional days. Neutral lipids (NL) were stained using Nile red and nuclei using DAPI. Neutral lipids labelling was quantified using Image J software (right). Scale bar: 50 µm (n= 10 to 13). Data are represented as mean +/- SEM, of a fold increase compared with the low glucose with OxLDL condition (E, F). Statistical analysis was performed using Mann-Whitney (C, D) and Friedman tests (E, F). *p<0.05, **p<0.01, ***p<0.001, ****p<0.0001

**Table 1:**
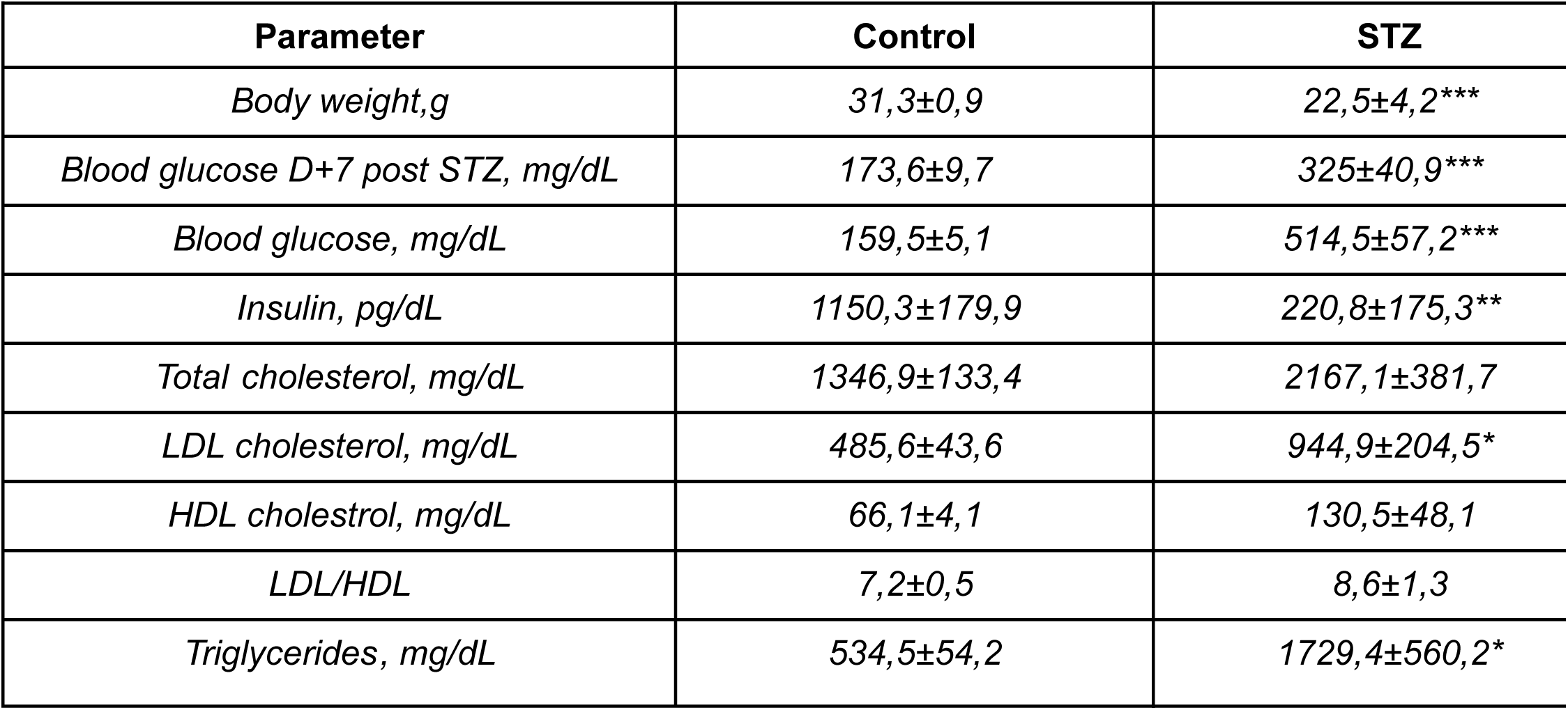
E**n**d **point plasmatic metabolic parameters of STZ and control LDLR^-/-^ mice.** Results are presented as the mean +/- SEM. Statistical analysis were performed using a Mann Whitney test (* p<0,05, ** p<0,01; *** p<0,001).

| **Parameter** | **Control** | **STZ** |
| --- | --- | --- |
| *Body weight,g* | 31,3±0,9 | 22,5±4,2*** |
| *Blood glucose D+7 post STZ, mg/dL* | 173,6±9,7 | 325±40,9*** |
| *Blood glucose, mg/dL* | 159,5±5,1 | 514,5±57,2*** |
| *Insulin, pg/dL* | 1150,3±179,9 | 220,8±175,3** |
| *Total cholesterol, mg/dL* | 1346,9±133,4 | 2167,1±381,7 |
| *LDL cholesterol, mg/dL* | 485,6±43,6 | 944,9±204,5* |
| *HDL cholestrol, mg/dL* | 66,1±4,1 | 130,5±48,1 |
| *LDL/HDL* | 7,2±0,5 | 8,6±1,3 |
| *Triglycerides, mg/dL* | 534,5±54,2 | 1729,4±560,2* |

As the formation of foam cells is associated with either altered OxLDL uptake and/or impaired cholesterol efflux ^16^, we analysed both mechanisms under elevated glucose concentration. Our analysis showed that after short exposures to OxLDL, no difference was observed in the intracellular accumulation of these lipoproteins in the presence of different concentrations of glucose, suggesting that SMC lipid accumulation is affected by high glucose concentration beyond the process of OxLDL uptake (Suppl. Fig. 1C). Consistently, SMCs cultured with a high glucose concentration demonstrated reduced free cholesterol efflux by more than 72 % and 35 % towards ApoA (a) and HDL (h) respectively compared to low glucose concentration (Suppl. Fig. 1D). Moreover, while the addition of OxLDL to the medium induced a significant increase in ATP-binding cassette A1 (ABCA1) and ATP-binding cassette G1 (ABCG1) transporter gene expression in SMCs, high glucose concentration did not affect this process (Suppl. Fig. 1E, F). Altogether, these results indicated that high glucose concentration leads to an impairment in lipid degradation by SMCs.

### High glucose concentration-induced lipid accumulation in SMCs is due to defective autophagy

Current literature suggests that lipid degradation occurs through an autophagy-related process in macrophages and SMCs ^17^. To investigate if elevated glucose concentrations could affect this process, we assessed LC3I/II expression in SMCs cultured in both glucose concentrations. Chronic incubation with high glucose concentration led to a 50 % decrease in LC3-II protein expression compared to low-rate glucose exposition and this was independent of a 3 day-treatment with OxLDL (Figure 2A and Supp. Fig. 2). Moreover, LC3-II expression in the presence of autophagy poison (Bafilomycin A), showed a dramatic decrease of the autophagy flux in the presence of high glucose concentration (Figure 2A, B). Analysis of the GFP-LC3- RFP-LC3ΔG probe evolution in these cells strengthened our observations as GFP/RFP ratio was significantly increased when cells were incubated with high glucose concentrations (Figure 2C).

**Figure 2:**
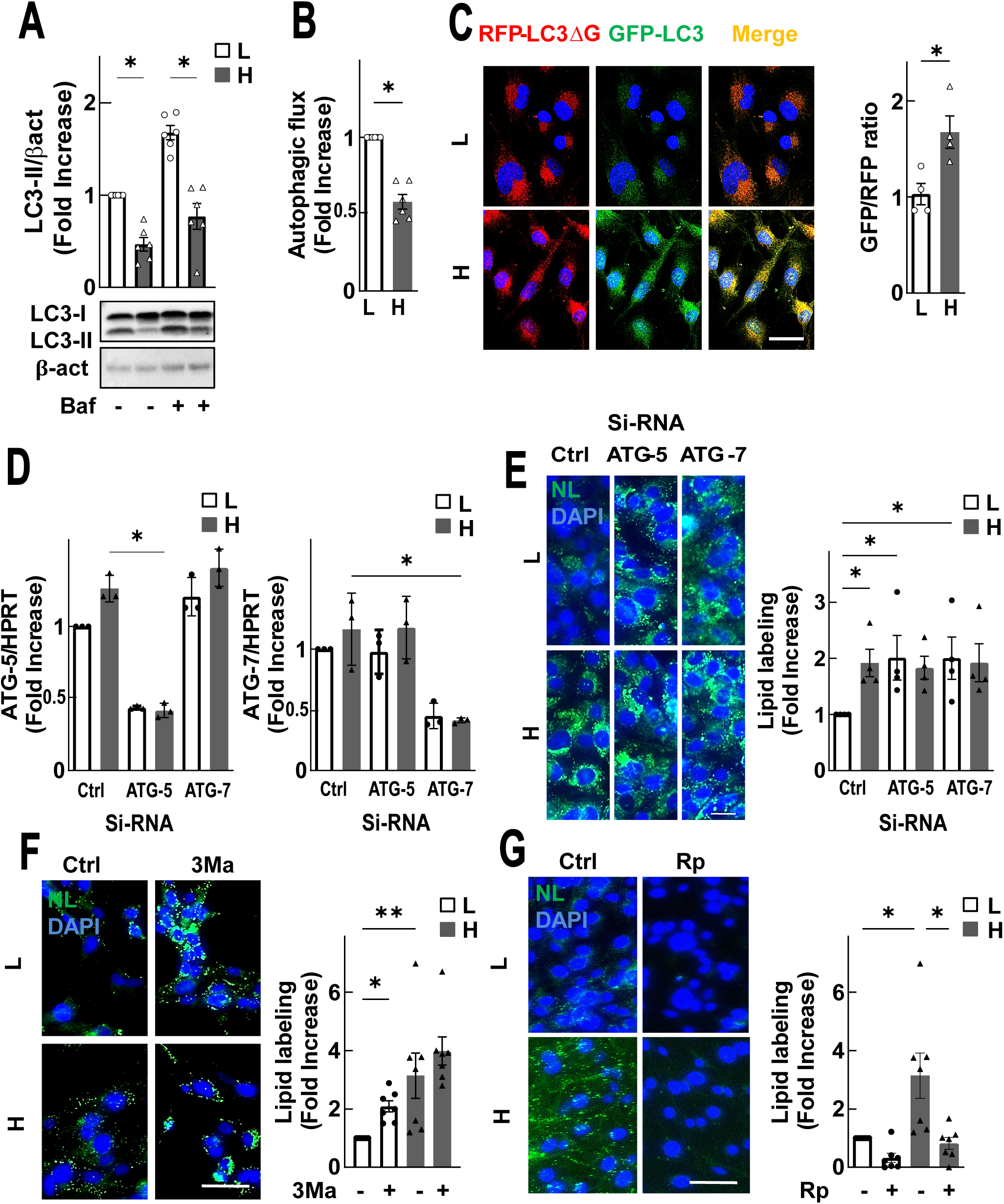
High glucose concentration induced lipid accumulation in SMCs is due to defective autophagy. SMCs were cultured in low (L, 1 g/L) or high (H, 5 g/L) glucose medium for 7 days A: Western Blot analysis of LC3 with or without bafilomycin (Baf,100 nM) 6 hours before the end of the experiment (n=5). B: Autophagic flux of SMCs treated as described in A (n=4). C: Representative confocal images (left) of low (L) and high (H) glucose concentration cultured hSMCs following transfection with the GFP-LC3-RFP-LC3ΔG plasmid as described in the Material and Methods. The GFP/RFP ratio (right), estimates the autophagic flux activity (n=4). Scale bar: 20 µm. D: Quantitative real-time PCR analysis of the mRNA levels of ATG-5 and ATG-7 (n=3). E-G: Foam cell formation in the presence of 200 µg/ml of oxLDL under various experimental conditions: E: Representative images of neutral lipids accumulation (left, green) of hSMCs transfected either with control (ctrl), ATG-5 or ATG-7 si-RNA and lipid labeling quantification (right) (n=4). F, G: Representative images of neutral lipid accumulation (left, green) of hSMCs treated with (+) or without (-) 3-methyladenine (3Ma, 20 µM, F) or rapamycin (Rp, 100 nM, G) and lipid labeling quantification (right) (n=6-7). DAPI was used to visualize the nuclei. Scale bars: 50 (C) and 70 µm (E-G). Results are presented as the mean +/- SEM, as a fold increase of the low glucose without the treatment (A-C, F, G) or low glucose with si-RNA control (D, E) conditions. Statistical analysis were performed using the Friedman test (A, D-G), Wilcoxon test (B), and unpaired t test (C). *p<0,05, **p<0,01.

To confirm the causal role of autophagy regulation in high glucose-induced foam SMC formation, we assessed the effects of genetical or pharmacological autophagy modulation in both low and high glucose conditions. First, ATG5 (autophagy related gene 5) and ATG7 (autophagy related gene 7) were knocked down by si-RNA to effectively abolish the autophagic process by limiting the expression of its partners (Figure 2D). ATG5 and ATG7 deficiency promoted foam cell formation in low glucose condition, without interfering with the effects observed when cells are cultured with high glucose concentration (L/ATG5 2.01±0.4, H/ATG5 1.8±0.2, *p<0.05, Figure 2E). These findings were further confirmed by the use of 3- methyladenine (3Ma), a specific autophagy inhibitor, that induced a two-fold increase in foam cell formation with low but not high glucose concentration (Figure 2F). A two-fold accumulation of total cholesterol in SMCs, especially cholesterol esters, and an 80 % decrease of cholesterol efflux towards ApoA were also observed in the presence of 3Ma in low glucose concentration (Supp. Fig. 3A-C). In contrast, the addition of the mTORC1 inhibitor rapamycin (Rp), a potent autophagy activator, significantly protected SMCs from high glucose-induced lipid accumulation, and attenuated foam cell formation (H/Rp(-) 3.1±0.5, H/Rp(+) 0.7±0.1, *p<0.05, Figure 2G). Consistent with these results, Rp treatment also limited the total cholesterol and cholesterol ester accumulation, along with the recovery of the cholesterol efflux to ApoA (Supp. Fig. 3D-F). Together, these findings confirmed that the downregulation of autophagy by high glucose is responsible for increased foam cell formation.

### PI3Kγ drives increased foam SMC formation under high glucose concentration

We aimed at deciphering the molecular mechanism underlying the effects of high glucose. The PI3K/Akt/mTOR pathway is largely involved in autophagy regulation in vascular foam SMCs ^18^. Using the pan-PI3K inhibitor LY294002 we confirmed that PI3K activity is involved in high glucose-induced foam SMC formation (Figure 3A). Nevertheless, the exact isoform of PI3K, involved in this process under high glucose conditions still needed to be identified. We therefore used specific PI3K isoform inhibitors against PI3Kα (A66), PI3Kβ (AZD-6482) and PI3Kγ (IPI-549) on SMCs cultured with low or high glucose concentrations. We found that the PI3Kγ inhibitor IPI-549 was the most effective at preventing foam SMC formation induced by high glucose (Figure 3B). Accordingly, in high glucose conditions, IPI- 549 markedly decreased intracellular total cholesterol content and increase free cholesterol generation (Figure 3C). In addition, IPI-549 restored 80 % of the efflux to ApoA in comparison to high glucose condition (Figure 3D). Finally, to confirm the involvement of PI3Kγ in these processes, we performed the same experiments in primary SMCs obtained from WT or PI3KγKO mice. In murine cells, genetic invalidation of PI3Kγ (Figure 3E) prevented high glucose-induced foam SMC formation. Altogether, these data identified PI3Kγ as a specific regulator of high glucose-dependent foam SMC formation.

**Figure 3:**
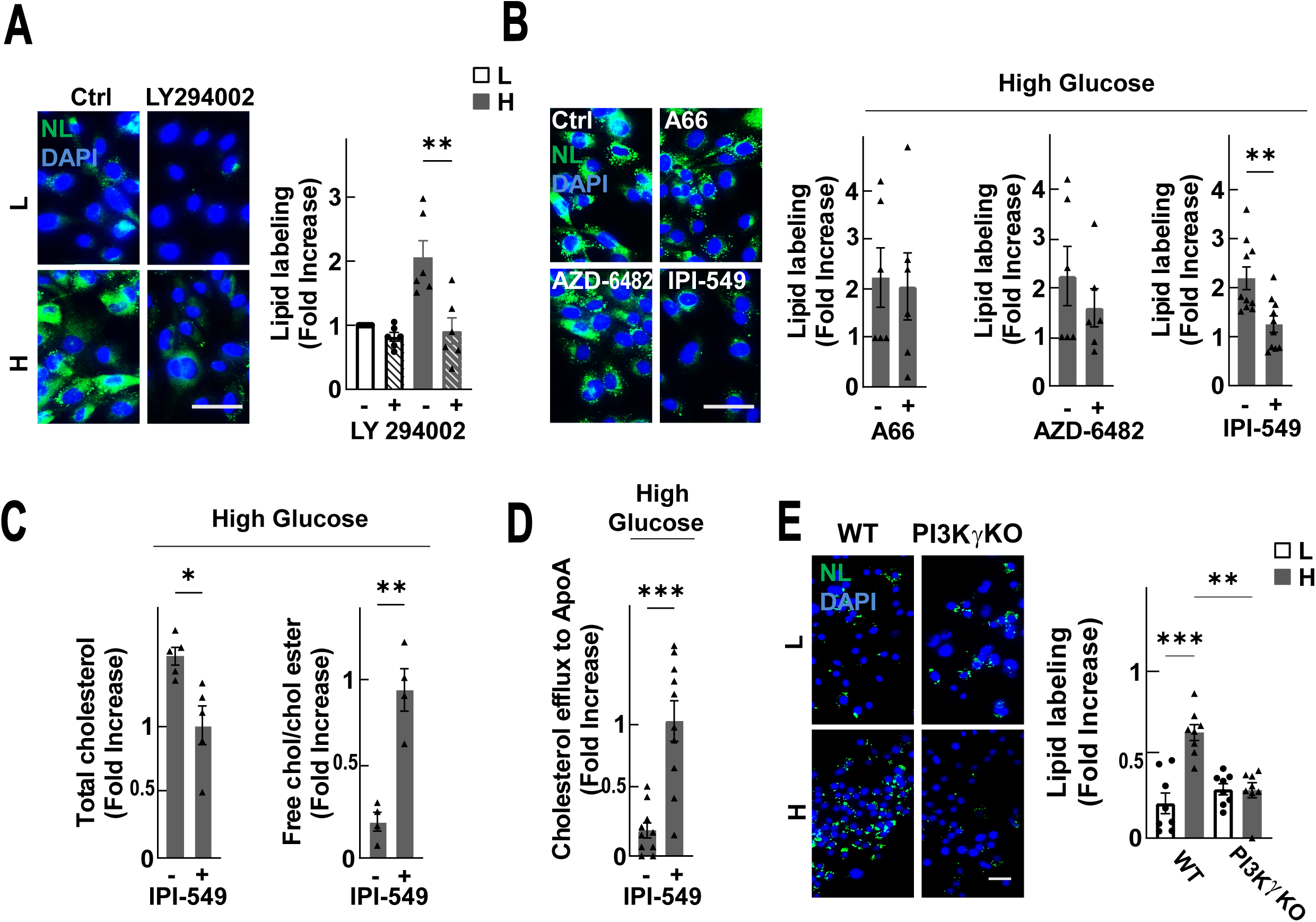
PI3Kγ drives increased foam SMC formation induced by high glucose concentration. hSMC were cultured with oxLDL (OxL, 200 µg/ml) in low (L) and high (H) glucose medium and then treated with the indicated treatment. A: Representative images of neutral lipids accumulation (left, green) and quantification (right) of hSMCs cultured with (+) or without (-) LY294002 (10 µM). DAPI was used to visualize nuclei. (n=6) B: Images of neutral lipids accumulation (left, green) and quantification (right) of hSMCs treated under class I PI3K isoforms inhibitors A66 (10 µM), AZD-6482 (10 µM) or IPI-549 (1 µM). DAPI was used to visualize nuclei. (n=6 to 10). C: Quantification of the total cholesterol level and free cholesterol/cholesterol ester ratio in hSMCs incubated in the presence (+) or absence (-) of IPI-549. (n= 4 to 5) D: Cholesterol efflux to ApoA assessed in hSMCs cultured as described in C. (n=10) E: Representative images of neutral lipids accumulation (left, green) and quantification (right) in wild-type (WT) or PI3Kγ knockout (PI3KγKO) mSMCs treated as described in Figure 1F. Nuclei were labeled using DAPI. (n=8) Scale bars: 50 µm. Results are presented as the mean +/- SEM and as a fold increase of the low glucose condition (A-D). Statistical analysis were performed using the Friedman test (A), unpaired t test (B-D), and Kruskal Wallis test (E). *p<0,05, **p<0,01, ***p<0,001.

### Blocking PI3Kγ reverses high glucose-induced autophagy alterations by modulating the nuclear translocation of TFEB

To further decipher how PI3Kγ inhibition counteracted foam SMC formation, we analysed autophagy dynamics in both human and murine SMCs (hSMC and mSMC). In hSMCs, pharmacological inhibition of PI3Kγ led to an increase in LC3 staining (Figure 4A) in high glucose condition. Moreover, the expression of autophagy-related genes, lysosomal- associated membrane protein 1 (LAMP-1 and LC3), reduced by 35% in high glucose condition, was restored by IPI-549 (Figure 4B). Consistently, genetic invalidation of PI3Kγ in mSMCs did not impact LC3 expression in high glucose compared to low glucose condition (Figure 4C). Autophagic flux assays were carried out on mSMCs, pre-treated with low or high glucose medium, using the GFP-LC3-RFP-LC3ΔG probe. As observed in Fig. 4D, the GFP/RFP ratio observed in high glucose is higher in WT SMCs in comparison to PI3Kγ-KO SMCs, meaning a higher autophagic flux when PI3Kγ is absent (Figure 4D). Moreover, high glucose significantly decreased LAMP-1 and LC3 gene expression by 40 % and 30 % respectively, in murine WT SMCs compared to low glucose medium (Figure 4E). Interestingly, the lack of PI3Kγ in high glucose condition maintained autophagy gene expression at the same levels as in low glucose condition (Figure 4F). Collectively, our data demonstrate that specific modulation of PI3Kγ activity in high glucose conditions is sufficient to restore the autophagic process in SMCs. To delineate the molecular mechanisms linking PI3Kγ to the regulation of autophagy, we investigated the involvement of TFEB, a master transcriptional regulator of autophagy: indeed, nuclear translocation of TFEB induces the transcription of key autophagy genes, and can be regulated through PI3K/mTOR pathway ^19^. Genetic knockdown of TFEB enhanced foam cell formation under low-glucose conditions without affecting high-glucose conditions despite an effective reduction in TFEB expression in both conditions (Figure 5A, B). Furthermore, TFEB downregulation led to decreased LAMP-1 and LC3 gene expression in cells cultured under low but not high glucose conditions, suggesting a key role of this transcription factor in high glucose-induced defective autophagy (Figure 5C, 5D). Western blot analysis revealed a significant decrease in TFEB nuclear translocation in SMCs cultured in high glucose compared with the low glucose condition (L: 1.98±0.28, H: 0.82±0.18; Figure 5E). Additionally, chronic exposition of SMCs to high glucose concentration induced the phosphorylation of TFEB on Serine 211, associated with a cytoplasmic retention of TFEB (Figure 5F). Incubation of SMCs with the PI3Kγ specific inhibitor IPI-549 significantly reduced S211-TFEB phosphorylation induced by high glucose (Figure 5F).

**Figure 4:**
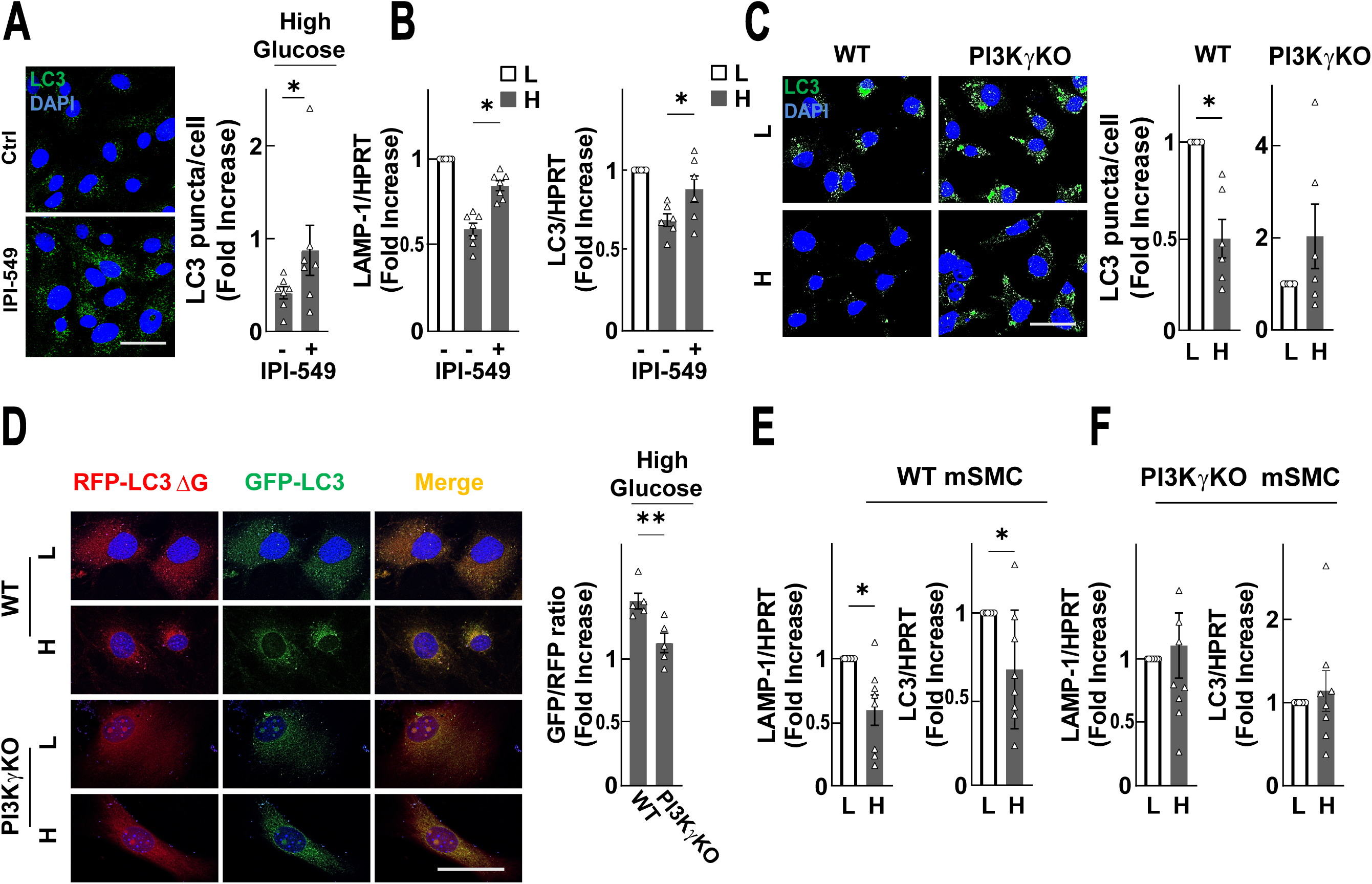
**Blocking PI3Kγ reverses high glucose-induced autophagy alterations**. A: Confocal images (left) of LC3 staining in hSMCs cultured in high glucose concentration treated with (+) or without (-) IPI-549 (1µM). LC3 puncta quantification (right) was performed using Image J software. (n=6) B: LAMP-1 and LC3 genic expression in hSMC stimulated with low (L) or high glucose (H) with (+) or without (-) IPI-549 (n=7). C: Confocal images (left) of LC3 staining in WT and PI3KγKO mSMCs cultured in low (L) or high glucose (H) medium. LC3 puncta quantification (right) was performed using Image J software. (n=6). D: Representative confocal images (left) of low (L) and high (H) glucose concentration cultured WT and PI3KγKO mSMCs following transfection with the GFP-LC3-RFP-LC3ΔG plasmid as described in the Material and Methods. Scale bar: 20 µm. The GFP/RFP ratio (right) was quantified using Image J software. E, F: LAMP-1 and LC3 genic expression in WT (E) and PI3Kγ KO (F) mSMCs stimulated with low (L) or high (H) glucose medium (n=7 to 8). Scale bars: 50 µm (A, C) and 20 µm (D). Results are presented as the mean +/- SEM, as a fold increase of low glucose condition. Statistical analysis were performed using Wilcoxon test (A, C-F) and Friedman test (B). *p<0,05, **p<0,01.

**Figure 5:**
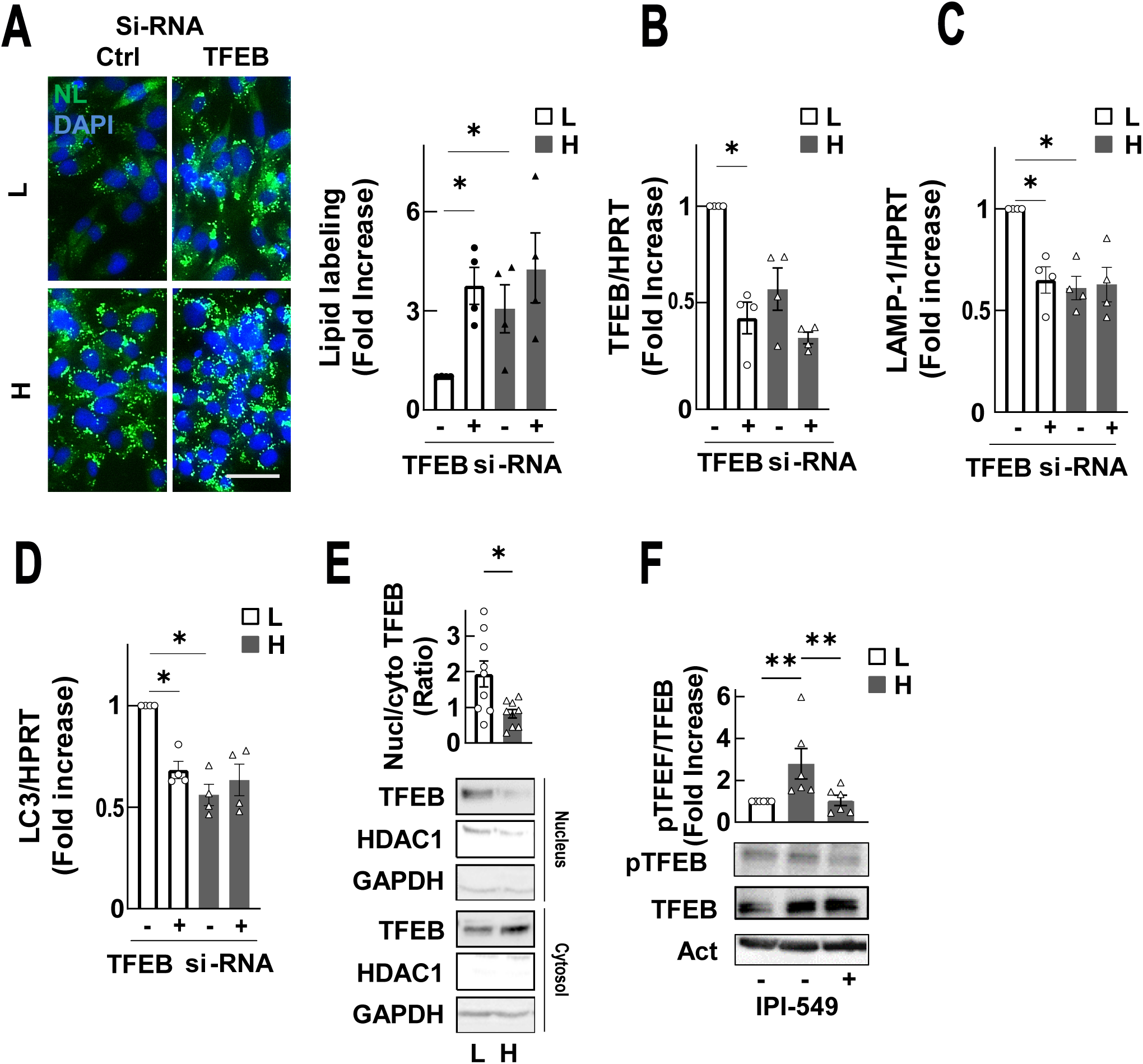
**PI3Kγ inhibition impaired high glucose-induced TFEB phosphorylation**. A: Representative images of neutral lipids accumulation (left, green) of hSMCs transfected with either control (ctrl) or TFEB si-RNA and lipid labelling quantification (right). Nuclei were visualized with DAPI staining. Scale bar: 50 µm. (n=4). B-D: Quantitative real-time PCR analysis of the mRNA levels of TFEB (B), LAMP-1 (C) or LC3 (D) (n=4). E,F: hSMCs were cultured in low (L, 1 g/L) or high (H, 5 g/L) glucose medium for 7 days. E: Ratio of TFEB protein expression in nuclear (Nucl) and cytoplasmic (Cyto) fractions, following Western blot analysis (n=9) F: Phospho-Ser211 and total TFEB protein expression with (+) or without (-) IPI-549 (1µM), following Western Blot analysis (n=6). Results are presented as the mean +/- SEM, as a fold increase of the low glucose with (A-D) or without (F) the si-RNA control condition. Statistical analysis were performed using the Friedman test (A-D, F) or t-Test (E). *p<0,05, **p<0,01.

Thus, the inhibition of PI3Kγ induces a desensitization of SMCs to high glucose- impaired autophagy, through the recovery of TFEB nuclear translocation, enabling the subsequent expression of autophagy key genes.

## DISCUSSION

Our results demonstrate that hyperglycemia promotes *in vivo* and *in vitro* SMC-derived foam cell formation. Using a wire injury-based model of accelerated atherosclerosis in hyperglycemic mice, we demonstrated that hyperglycemia led to lesions enriched in foam SMCs compared to normoglycemic mice. Previous studies in the literature demonstrated the impact of hyperglycemia on the acceleration of atherosclerotic plaque formation in the aortic sinus using ApoE^-/-^ mice rendered diabetic with streptozotocin ^20^. However, the role of SMCs in this context remained unclear. Our model, which uses a guidewire to denude the endothelium, has the peculiarity of a rapidly developing atherosclerosis characterized by a massive lipid infiltration in SMA-positive SMCs. This model has allowed us to demonstrate for the first time that elevated blood glucose levels have a deleterious effect on SMCs under atheromatous conditions by promoting their intracellular lipid accumulation.

*In vitro*, SMCs chronically exposed to high glucose medium in the presence of oxLDL significantly increase neutral lipid engulfment in comparison with low glucose condition. Our results, unlike the work of Xue and al., demonstrate that high glucose concentration did not induce any modification of LDL uptake nor decrease of cholesterol efflux receptor expression^16^. Indeed, while they reported a reduction of ABCG1 expression in SMCs cultured in oxLDL- and high glucose-enriched medium, our work rather shows that high glucose alone induces an alteration in autophagy. In the literature, lipid-associated autophagy is referred to as lipophagy ^21^. Our results therefore suggest that high glucose concentrations are capable of modifying this specific lipophagy mechanism.

In our study, high glucose concentration-impaired autophagy is due to the alteration of autophagy key gene expression. Interestingly, these detrimental effects have been found in the vascular wall from gestational and type 2 diabetes mellitus patients, supporting a defective autophagic activation in diabetic conditions ^22–24^. Accordingly, we found that elevated glucose concentrations reduced autophagy process through TFEB nuclear translocation impairment, reinforcing the role of TFEB in autophagy regulation in SMCs, as it has been proposed by Chen and al. ^25^.

Foam SMC formation has been associated with the PI3K class I pathway ^18^. Here we demonstrate an unexpected role of the gamma isoform of the PI3K family in response to high glucose in SMC foam cells. In foam macrophages, the role of the gamma isoform has been suggested but not clearly demonstrated ^26,27^. Chang and al. showed that in PI3Kγ-deficient mice, foam macrophage formation is reduced. This phenomenon likely originates from an indirect inhibitory effect on inflammation, resulting in a decreased foam cell content inside the plaque ^28^. Here, we identify for the first time that in high glucose concentration, PI3Kγ controls TFEB phosphorylation leading to decreased autophagy and lipid accumulation in SMCs. These data suggest that elevated glucose concentrations potentiate PI3Kγ activity. One possible mechanism could be a modification in GPCR expression or activation, as it has been proposed for P2Y12 expression in platelets of diabetic patients ^29^. However, the specific mechanism linking high glucose concentration to PI3Kγ activation needs further investigation.

Despite improvements in cardiovascular disease management, diabetic patients still present a higher risk of major thrombotic events and their recurrence compared to non-diabetic patients ^30,31^. Glycemic control has long been considered as the main factor capable of limiting those risks ^32,33^. Collectively, our work brings the evidence that hyperglycemia aggravates foam SMC formation through defective autophagy. Our results, together with data from literature, supporting the fact that hypoglycemic drugs could exert atheroprotective effects by promoting autophagy in SMCs, clearly demonstrate the importance of regulating this process in SMCs of diabetic patients, to avoid macrovascular complications ^23,24^. Moreover, our results shed light on the possibility of using of PI3K gamma inhibitors, which have passed safety clinical trials for the treatment of solid tumors, to prevent major cardiovascular events and potential recurrences in diabetic patients ^34^.

## Acknowledgments

We thank the Phenotyping Department of the UMS006 CREFRE-Anexplo Platform and the GeT-Santé facility (I2MC, Inserm, Génome et Transcriptome, GenoToul, Toulouse, France) and TRI-Genotoul (light microscopy; member of the national infrastructure France-Bioimaging Infrastructure supported by the French National Research Agency) for their technical support.

## Funding

This work was funded in part by Inserm, Foundation for medical research (FRM, Target PI3K) and French Foundation (FDF cardiovascular: SMC and Diabetes), and the FEDER- région Occitanie program (INSPIRE). H.L. and I.M. were supported by a research grant from the University of Toulouse.

## Data Availability Statement

Data will be made available upon reasonable request.

## Conflicts of Interest

The authors declare no conflict of interest.

## Notes

### Competing Interest Statement

The authors have declared no competing interest.

### Summary of Updates

We present an improved version of the abstract and the material and method sections. Most figures were also improved after several rounds of reviewing processes.

