## supplemental figures for "Hyperglycemia worsens smooth muscle foam cell formation through PI3Kγ-dependent defective autophagy"

*, ** equally contributed

### Supplementary Figures

**Supplementary Figure 1: High glucose concentration does not alter oxidized LDL uptake but leads to a decrease in cholesterol efflux.** A, B: Total cholesterol level (A) and free cholesterol/cholesterol ester ratio (B) were evaluated in hSMCs incubated for 7 days in either low (L, 1 g/L) or high (H, 5 g/L) glucose medium with or without OxLDL (OxL, 200 µg/mL) as described in material and method section (n=7). C: Representative images (left) of hSMCs cultured in low (L) or high (H) glucose medium during 7 days, supplemented with DiI labeled-OxLDL (Di-OxL) during 0 or 24 additional hours, and fluorimetric quantification (right) (n=4). Scale bar: 50 µm. D: Cholesterol efflux to ApoA (a) or HDL (h) in hSMCs (n=11). E, F: Quantitative real-time PCR analysis of the mRNA levels of ABCA1 (E) or ABCG1 (F) transporters in hSMCs (n=6). All experiments are represented as the mean +/- SEM and a fold increase of low glucose with (A, B) or without (E, F) OxLDL or to low glucose efflux to ApoA (D) conditions. Statistical analysis was performed using Friedman test (A, B, D-F) or the one-way ANOVA test (C). *$p<0.05$, **$p<0.01$.

**Supplementary Figure 2: High glucose concentration impairs autophagy with or without oxidized LDL.** A: LC3 protein expression analysis by Western Blot in SMC cultured with (+) or without (-) OxLDL (OxL, n=4). B: Confocal images (left) of LC3 staining. DAPI was used to visualize the nuclei. LC3 puncta quantification (right) was performed using Image J software (n=5). Scale bar: 20 µm. Results are presented as the mean +/- SEM and as a fold increase of the low glucose condition. Statistical analysis were performed using the Friedman test. *p<0,05, **p<0,01.

**Supplementary Figure 3: Total cholesterol, free cholesterol and cholesterol efflux are regulated by autophagy.** hSMC were cultured in oxLDL low (L) and high (H) glucose medium and then treated with the indicated treatment. A, B: Total cholesterol level (A) and free cholesterol/esterified cholesterol ratio (B) quantifications in hSMCs cultured with (+) or without (-) 3-methyladenine (3Ma, 20 µM). (n=4 to 6). B: Cholesterol efflux to ApoA was quantified in hSMCs cultured as described in A. (n=6). D, E: Total cholesterol level (D) and free cholesterol/esterified cholesterol ratio (E) quantification in hSMCs cultured with (+) or without (-) rapamycin (Rp, 100 nM) (n=4). F: Cholesterol efflux to ApoA was quantified in hSMCs cultured as described in D (n=5). Results are presented as the mean +/- SEM and a fold increase of the low glucose without treatment condition. Statistical analysis was performed using the Friedman test. *p<0,05, **p<0,01.

**Supplementary Figure 1: High glucose concentration does not alter oxidized LDL uptake but leads to a decrease in cholesterol efflux**

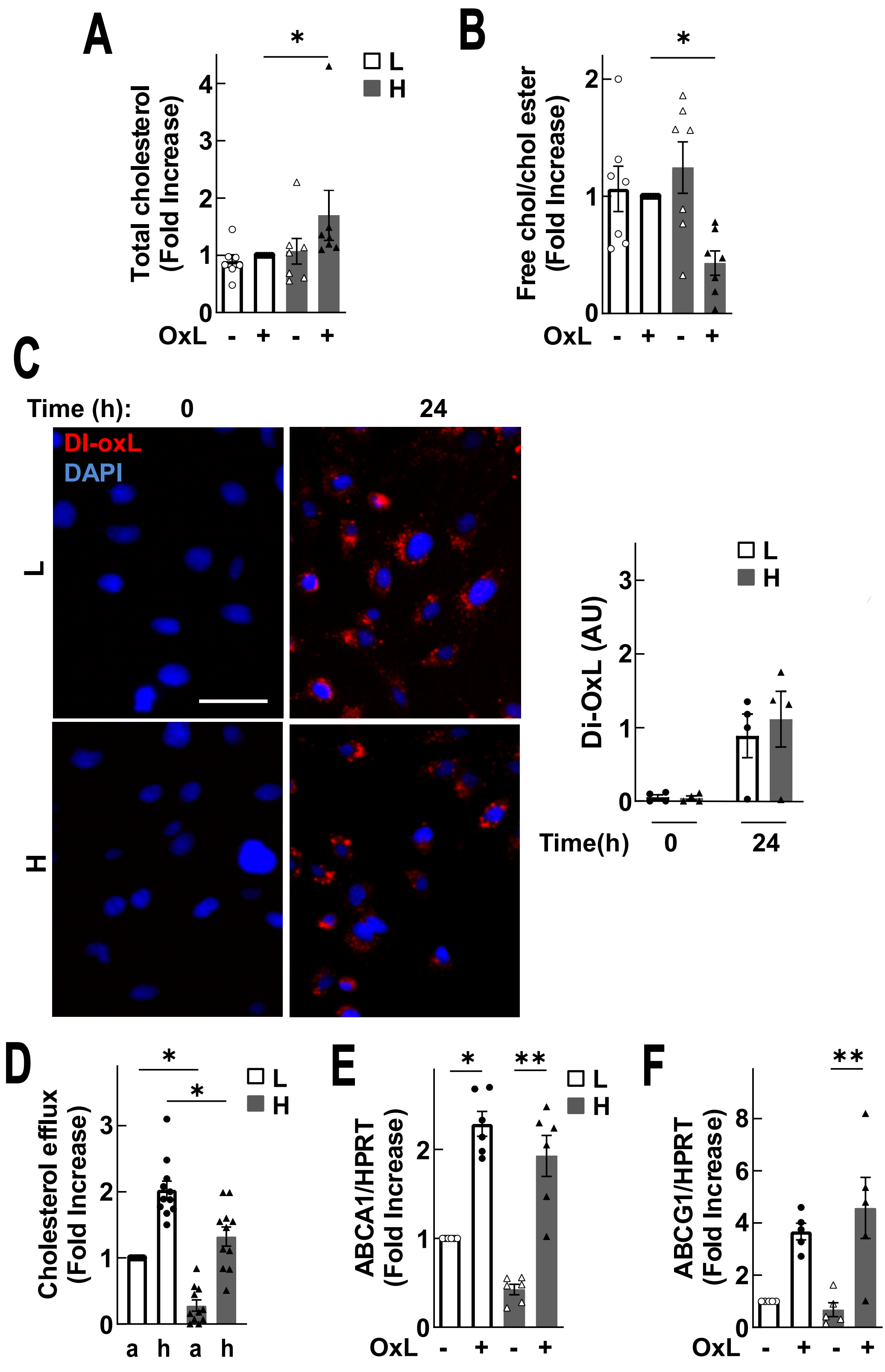

**Supplementary Figure 2: High glucose concentration impairs autophagy with or without oxidized LDL**

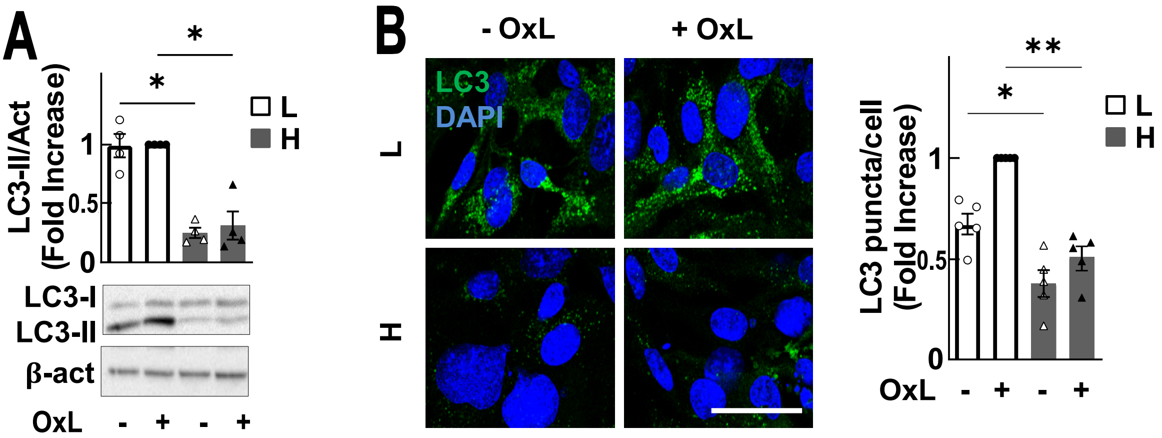

**Supplementary Figure 3: Total cholesterol, free cholesterol and cholesterol efflux are regulated by autophagy**

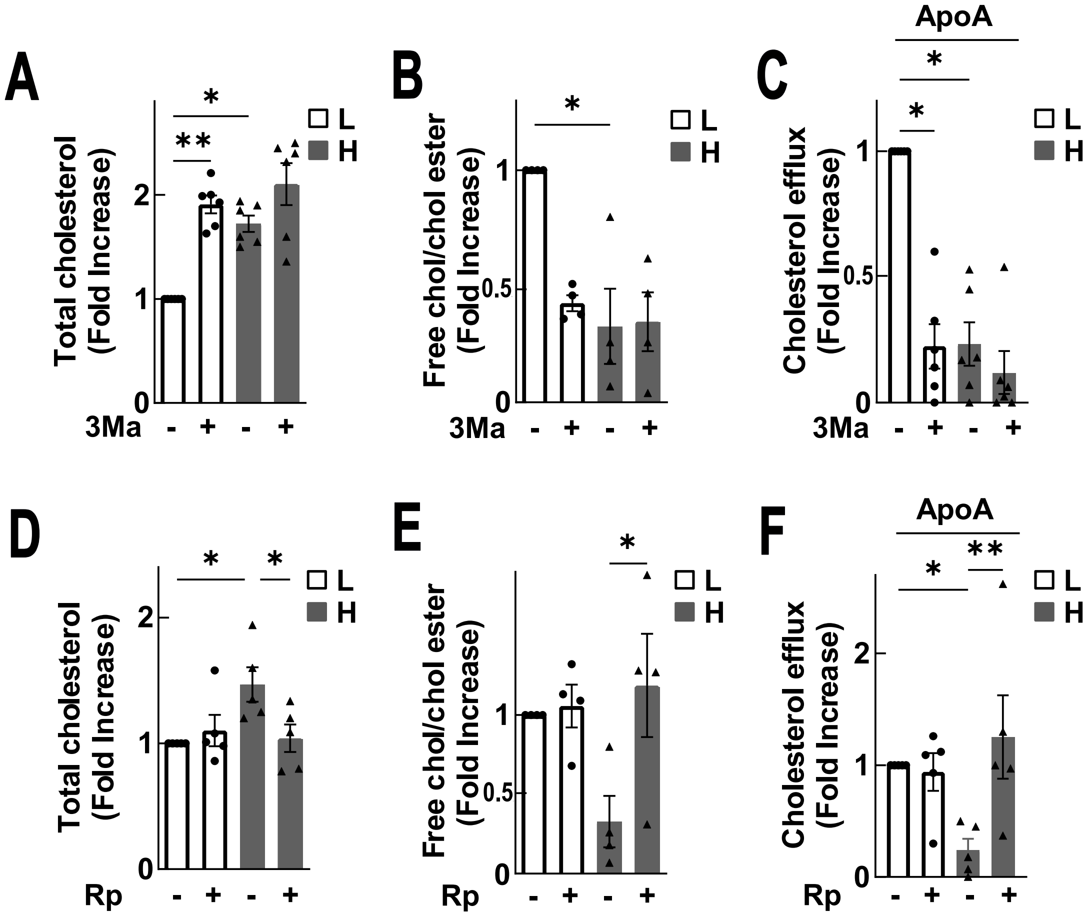
